# qRT-PCR/High Resolution Melting Analyses detected polymorphisms of *HSP-70* gene in Zebu cattle

**DOI:** 10.1101/483792

**Authors:** G.O. Onasanya, M. Okpeku, K.A. Thiruvekander, C. Sreekumar, G.K. Tirumurugaan, G.M. Msalya, T. M. S anni, O. Olowofeso, A.O. Fafiolu, M.A. A deleke, C.O.N. Ikeobi

## Abstract

Concerns over environmental heat stress have increased in recent years with the realization of global warming effects on environment and subsequently on animal production. Heat stress indicators help to monitor adaptability and survivability of livestock to thermal assault; a consequent effect of global warming. Heat shock proteins (HSPs) are highly conserved and are classified into six groups. Heat shock protein *(HSP)* 70 gene is a member of molecular chaperone, known to be highly expressed under stressful environmental and physiological conditions. In this study, we used a one-step qReal-Time PCR, followed by high-resolution melting analyses (HRMA) for detection and identification of polymorphism of *HSP-70* gene in four extant cattle breeds of (White Fulani, Sokoto Gudali, Red Bororo and Ambala). qRT-PCR/HRMA-based assay detected twelve (12) genetic variant groups of *HSP-70* gene. Ambala breed of Nigerian Zebu cattle had the highest number (7) of genetic variants of *HSP-70* gene followed by Red Bororo breed (6 genetic variants), Sokoto Gudali breed had 5 genetic variants while White Fulani breed had the least (4 genetic variants) number of genetic variants in the population of Nigerian Zebu cattle breeds examined. We therefore hypothesize that the detected variant groups can be interrogated for their possible effects on thermo-tolerance performance of zebu cattle bred under thermal assaults of tropical conditions.

## Introduction

Cattle industry is a very important sector of livestock husbandry in many developing countries all over the world. Cattle serve as valuable source of animal proteins and hide, and contribute in no small measure to national economy. Heat stress is a significant issue in the livestock industry.

The impact of heat stress on livestock, particularly on cattle can be overwhelming [1], resulting in reduced feed intake, reduced fertility, increased respiratory and heart rates, increased peripheral blood flow and profuse sweating, reduced milk production and lower milk quality [2]. These impacts according [2], can result in significant loss of income and increase in management costs. Losses to the cattle industry due to heat stress was estimated at about $900 million per annum [3]. The issue is more exacerbated with increased global warming, as such, the impact of heat stress needs to be addressed and ameliorated to maintain animal health status, adaptability, survivability and performance.

Heat shock proteins (*HSPs*), are a group of proteins conserved in both prokaryotic and eukaryotic organisms. The expression of HSP genes are related to thermal stress, they play vital roles in cell response to environmental stress [4, 5]. Studies have shown that the expression of HSP genes affects follicular development, embryonic survival and pregnancy maintenance [6-8]. For this reason, the functional characterization of this gene family is important. The *HSP*-70 protein, also referred to as *HSP*-70 gene is a member of the *HSP* family, located on 23q13 in the bovine genome [9]. It is a member of a molecular chaperone implicated to be highly expressed under stressful environmental and physiological conditions [10].

*HSP*-70 gene as a member of the *HSPs* sub-family has been found to play essential roles in cellular protection, immune response, protein synthesis, cyto-skeletal protection, protein folding and unfolding, protein translocation and regulation of steroid hormone receptors, transportation, re-folding of protein, protecting proteins from cellular stress, inhibitory apoptosis and adaptation during and after thermal assault [11-13]. In addition to its functional importance, its close proximity to the major histocompatibility complex genes suggests that the *HSP*-70 gene as a powerful candidate marker for health, reproduction and productive traits [14]. For these reasons, studies of polymorphisms in this gene and resultant phenotypic traits in both livestock and human beings have increased [13, 15-17].

Detection of genetic variants in *HSP-70* gene can potentially be exploited for identification of animals tolerant/resistant to thermal stress through thermo-tolerance traits association studies so as to drift herds toward superior thermo-tolerant ability; through improvement of heat vulnerable stocks with thermo-tolerant stocks. Also, the identified genetic variants could be important for developing and managing livestock in the face of climate change, more efficient resource utilization and improved production performance in terms of fertility, milk production, feed intake, growth, conception rates and animal health [12, 13].

High Resolution Melting (HRM) is a post-PCR analysis technique, capable of discrimination among DNA sequences based on their composition, length, GC content, or strand complementarity [18]. It is a simple and fast technique based on PCR melting (dissociation) curve techniques, enabled through improved double-stranded DNA (dsDNA)– binding dyes along with next-generation real-time PCR instrumentation and analysis software [19]. The combine usage of High Resolution Melting analysis (HRMA) with quantitative real time polymerase chain reaction (qRT-PCR) permits efficient and rapid genotyping and detection of polymorphism in double stranded DNA (Bester et al., 2012; Gori et al., 2012; Yang et al., 2016) [20-22]. This technique generates DNA melt curve profiles that are very specific and sensitive enough to distinguish sequence variation enabling mutation scan, methylation and genotyping analysis. In the last few years, an increasing number of research articles have been published on qReal-Time PCR /HRMA-based assay, especially in the identification of human and animal pathogens [23-25]. The evolution of qRT-PCR/HRMA based assay involving the use of fluorescent intercalating dye (SYBR Green) with a new generation of light cyclers is a high throughput technology with an established average temperature range of 66 °C to 96 °C for mutation analyses, genetic variants analyses, polymorphism study and SNP genotyping of large population which has become emerging cutting edge technology with high efficiency and cost-effectiveness [26, 27]. This technology is gradually becoming a technique of choice for rapid genotyping and study of polymorphism [20-22].

Previous studies in Zebu cattle breeds suggested that they are superior in adapting to tropical climatic conditions compared to their European counterparts [28]. Zebu cattle are naturally adapted to a wide range of agro-climatic conditions [12, 13]. Nigeria is a tropical country with severe influence of thermal stress that significantly affects production performance of livestock. Nigerian indigenous cattle breeds are predominantly Zebu cattle. They are particularly adapted to the Northern part of the country with the highest heat assault. However, very little effort has been made to understand their heat stress status.

We therefore, evaluated polymorphism of *HSP*-70 gene in indigenous Zebu cattle, using qRT-PCR/HRMA-based assay in an attempt to characterize and gain understanding of genetic variation and polymorphism of *HSP*-70 gene in Nigerian zebu cattle breeds.

## Materials and methods

### Sampling regions and experimental animals

A total of ninety (90) adult bulls from across four extant breeds of Nigerian Zebu cattle comprising of White Fulani (25), Sokoto Gudali (21), Red Bororo (21) and Ambala (23) were sampled across ten (10) states in the Northern parts of Nigeria. Informed decision was obtained from farmers and herd owners before animal inclusion in the study. The animals originated from different herds and were reared under the traditional extensive system where they grazed during the day on natural pasture containing forages such as Stylo (*Stylosanthes gracilis*), Leucaena (*Leucaena leucocephala*) and Guinea grass (*Panicum maximum*), crop residues and scavenged on kitchen wastes whenever available.

### Sample collection for genomic DNA isolation

From each of the 90 samples, 200 g of skin tissue sample was excised from each animal prior to bleeding in the abattoir/slaughter house and same were sliced into less than 0.5 cm (about 1 g weight) and quickly submerged into 0.5 ml Eppendorf tubes containing RNA*later* reagent. Subsequently, the samples were transported on ice to laboratory and stored at −20°C until further analyses.

## DNA extraction

The protocol for DNA extraction employed for this study was according to HiPurA™ Multi-Sample DNA Purification procedure, MolBio™ Himedia^®^, Mumbai, India. 25 mg of skin tissue was fetched from each of the 90 samples of skin tissue preserved in RNA*later.* Samples were weighed using sensitive scale, then shredded into smaller pieces and transferred into 2 ml collection tube. 180 μl of resuspension buffer was added into the 2 ml collection tube containing the shredded skin tissue. Thereafter, 20 μl of proteinase K solution was added and thoroughly mixed by vortexing for proper tissue digestion. Incubation of the samples was done using ACCUBLOCK™ digital dry bath at 55 °C for 2-4 hours until the tissue was completely digested with no residues. During incubation, the samples were mixed occasionally by vortexing. After digestion, the samples were vortexed briefly for 30 seconds.

The preparation of lysate: lysation was done by adding 200 μl of lysis solution to 2 ml collection tube containing digested tissue, vortexing thoroughly for 15 seconds and subsequently incubated using ACCUBLOCK™digital dry bath at 70 °C for 10 minute to generate lysate. To prepare lysate for binding to the spin column, 200 μl of ethanol (100%) was added to the lysate obtained and mixed thoroughly by gentle pipetting and transferred into HiElute Miniprep spin column and centrifuged at 10000 rpm for 1 minute using Thermo Scientific Nanofuge (MCROCL 21/21R) micro-centrifuge. The flow–through liquid was discarded and the column was placed into a fresh 2 ml collection tube. 500 ul of dilute pre-wash solution (12 ml pre-wash + 18 ml ethanol) was added to the column containing the lysate and was centrifuged at 10, 000 rpm for 1 minute.

The flow-through liquid was subsequently discarded. In a separate step, 500 μl of diluted wash solution (8 ml of wash solution + 24 ml of ethanol) was added to the column containing the lysate and was centrifuged at 13,000 rpm for 3 minutes and the flow through liquid was discarded. The column was further centrifuged at 13, 000 rpm for 1 minute using and the collection tube containing flow–through liquid was discarded. The column was then placed into a new 2 ml collection tube and 100 ul of elution buffer added directly into the column and incubated for 5 minutes at room temperature (15-25 °C) and then centrifuged at 10, 000 rpm for 1 minute to elute DNA into collection tube after which the column was discarded. Eluted DNA was briefly incubated at 60 °C to free the DNA of any contamination and was subsequently stored at −20 °C for further analyses. Quality and quantity of DNA was estimated using Thermo Scientific-NanoDrop 2000 spectrophotometer (Shimadzu co-operation, Japan).

### Primer sequence and target regions

The *HSP-70* gene primers set (Table 1) used for this study was obtained from the earlier works of Bhat *et al.* (2016) and were optimised for primer specificity. The fragment size for *HSP-70* gene was 295 bp covering coding region in exon 1.

**Table 1.**
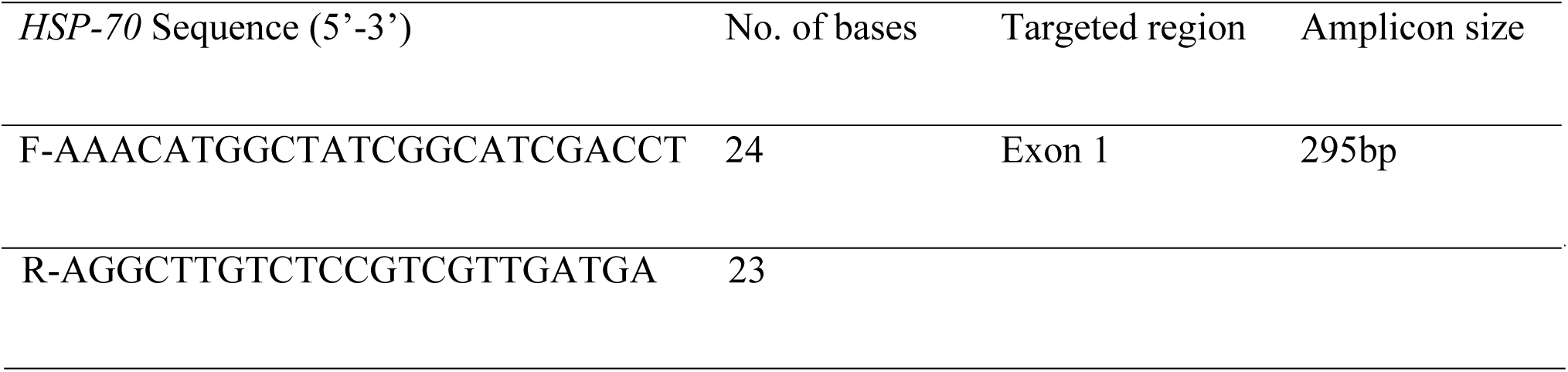
Heat Shock protein (HSP) 70 gene primer sequence and target region (181…475 bp)

### Polymerase chain reaction and amplification condition

The PCR reactions were carried out in a total volume of 15 μl containing template DNA of 1.0 μl, 1.0 μl of each of forward and reverse primers, 7.5 μl PCR Master Mix (2x) (GeNei™ Red Dye PCR Master Mix) and 4.5 μl of nuclease free water. PCR amplification was performed in a TaKaRa Thermal Cycler Dice™ version III (Takara Bio Inc., Japan). The amplification condition involved initial denaturation at 94 °C for 5 minutes, followed by 45 cycles of denaturation at 94 °C for 60 seconds, annealing temperature of 65°C for 45 seconds, extension at 72 °C for 1 minute followed by final extension at 72 °C for 7 minutes. PCR products were evaluated using 2% agarose gel electrophoresis after staining with 1 ug/ml of ethidium bromide and visualized under Bio-RAD Gel Doc™ XR+ Imaging System version 5.1 (Gel Documentation Molecular Imager, Bio-Rad Laboratories, Inc., U.S.A.).

### qRT-PCR and HRMA based assay

Heat shock protein 70 amplified fragments were purified prior to qRT-PCR procedure. 20 μl PCR products of *HSP* 70 gene were detected by gel electrophoresis on 2% agarose gel stained with ethidium bromide and viewed under Bio-RAD Gel Doc™ XR+ Imaging System version 5.1 (Gel Documentation Molecular Imager, Bio-Rad Laboratories, Inc., U.S.A.), the gel bands were carefully excised and placed into sterilised vials and stored at −20°C. The excised gel bands were then manually digested and centrifuged for 5 minutes at 10,500 rpm. A layer of supernatant (purified DNA) was formed on the surface of the gel, the purified supernatant (DNA fragment) was treated as DNA template and was subsequently used for qRT-PCR/ HRMA-based assay

Quantitative Real-time PCR followed by HRMA was carried out for identification and detection of possible genetic variants of *HSP-70* gene in four Nigerian Zebu cattle breeds. The qRT-PCR reactions were carried out on a total volume of 20 μl containing purified DNA fragment (template DNA) 1.0 μl, 1.0 μl of each of forward and reverse primers, SYBR green Master Mix (2x) of 10.0 μl, and 7.0 μl of nuclease free water. The qRT-PCR amplification was performed in a Roche LightCycler^®^96 with software version 1.01.01.0050 (Roche Diagnostics, Mannheim, Germany). The HRMA programme and qRT-PCR amplification condition consisted of pre-incubation for 5 minutes, followed by 45 cycles of denaturation at 95 °C for 10 seconds, annealing at 65 °C for 10 seconds and extension at 72 °C for 10 seconds. HRMA was performed by first heating to 95 °C for 1 minutes, cooling to 37 °C for 30 seconds, heating to 65 °C for 1 second and then melting with continuous acquisition (15 readings / °C) of florescence signal up to 97°C. Genetic variants of *HSP-70* gene generated across breeds of Nigerian *Bos indicus* were analysed using Roche High Resolution Melting Software version 1.1.0.1320. Individual samples of *HSP-70* gene were differentiated into distinct genetic variants via high resolution melting curve profiles (derivative HRM curve / dissociation curve, differential plot and normalised melt curves) as depicted with distinct SYBR green (dye) fluorescence: Purple (PRP), Red (RED), Orange (ORG), Green (GRN), Lemon (LMN), Brown (BRN), Chocolate (CHO), Yellow (YLO), Magenta (MGT), Blue (BLU), Army green (AGN) and Navy blue (NBL) where individual fluorescence depicted distinct genetic variants [21, 22].

### Quantitative Real-Time polymerase chain reaction / amplification condition and high resolution melting analyses-based assay programme

Quantitative Real-time PCR followed by HRMA was carried out for identification and detection of a possible genetic variant of *HSP-70* gene in four Nigerian cattle breeds. The qRT-PCR reactions were carried out on a total volume of 20 μl containing purified DNA fragment (template DNA) 1.0 μl, 1.0 μl of each of forward and reverse primers, SYBR green Master Mix (2x) of 10.0 μl, and 7.0 μl of nuclease free water. The qRT-PCR amplification was performed in a Roche LightCycler^®^96 with software version 1.01.01.0050 (Roche Diagnostics, Mannheim, Germany). The HRMA programme and qRT-PCR / amplification condition consisted of pre-incubation for 5 minutes, followed by 45 cycles of denaturation at 95 °C for 10 seconds, annealing at 65 °C for 10 seconds and extension at 72 °C for 10 seconds. HRMA was performed by first heating to 95 °C for 1 minutes, cooling to 37 °C for 30 seconds, heating to 65 °C for 1 second and then melting with continuous acquisition (15 readings / °C) of florescence signal until 97°C.

Possible genetic variants of *HSP-70* gene were evaluated via qReal-Time PCR followed by HRMA-based assay in four Zebu breeds of Nigerian cattle [20-22]. Subsequently, qRT-PCR / HRM data generated from across the breeds of Nigerian *Bos indicus* for *HSP* 70 gene were analysed using Roche High Resolution Melting Software version 1.1.0.1320. Individual samples of *HSP-70* gene were differentiated into distinct genetic variants via high resolution melting curve profiles (derivative HRM curve / dissociation curve, differential plot and normalised melt curves) as depicted with distinct SYBR green (dye) fluorescence; Purple (PRP), Red (RED), Orange (ORG), Green (GRN), Lemon (LMN), Brown (BRN), Chocolate (CHO), Yellow (YLO), Magenta (MGT), Blue (BLU), Army green (AGN) and Navy blue (NBL) where individual fluorescence depicted distinct genetic variants [21, 22].

## Results and Discussion

Detection of genetic variants in *HSP-70* gene in four Nigerian Zebu cattle breeds was studied using qRT-PCR/HRMA-based assay. The amplified gene spanned a coding region of approximately 295 bp. Quality of the amplified region was tested using the normalized qRT-PCR and the normalized HRMA techniques separately and then the equipment software was recalibrated to combine qRT-PCR with HRMA.

We detected dissociation of the *HSP-70* gene sequence at delta TM discrimination/dissociation (50%) and curve shape discrimination (50%), and fluorescence normalization sensitivity at pre-melting temperature (73.766 °C to 74.766 °C) and post melting temperature range was 94.966 °C to 95.966 °C (Fig. 1 and 2). Each dissociation produced three (3) distinct florescence that we tagged as wild type, homozygous and the heterozygous alleles of *HSP-70* gene respectively. Differential plot curve of the gene variants were also observed when the software associated with the equipment was primed to combine qRT-PCR with HRMA (Fig. 3).

**Fig 1.**
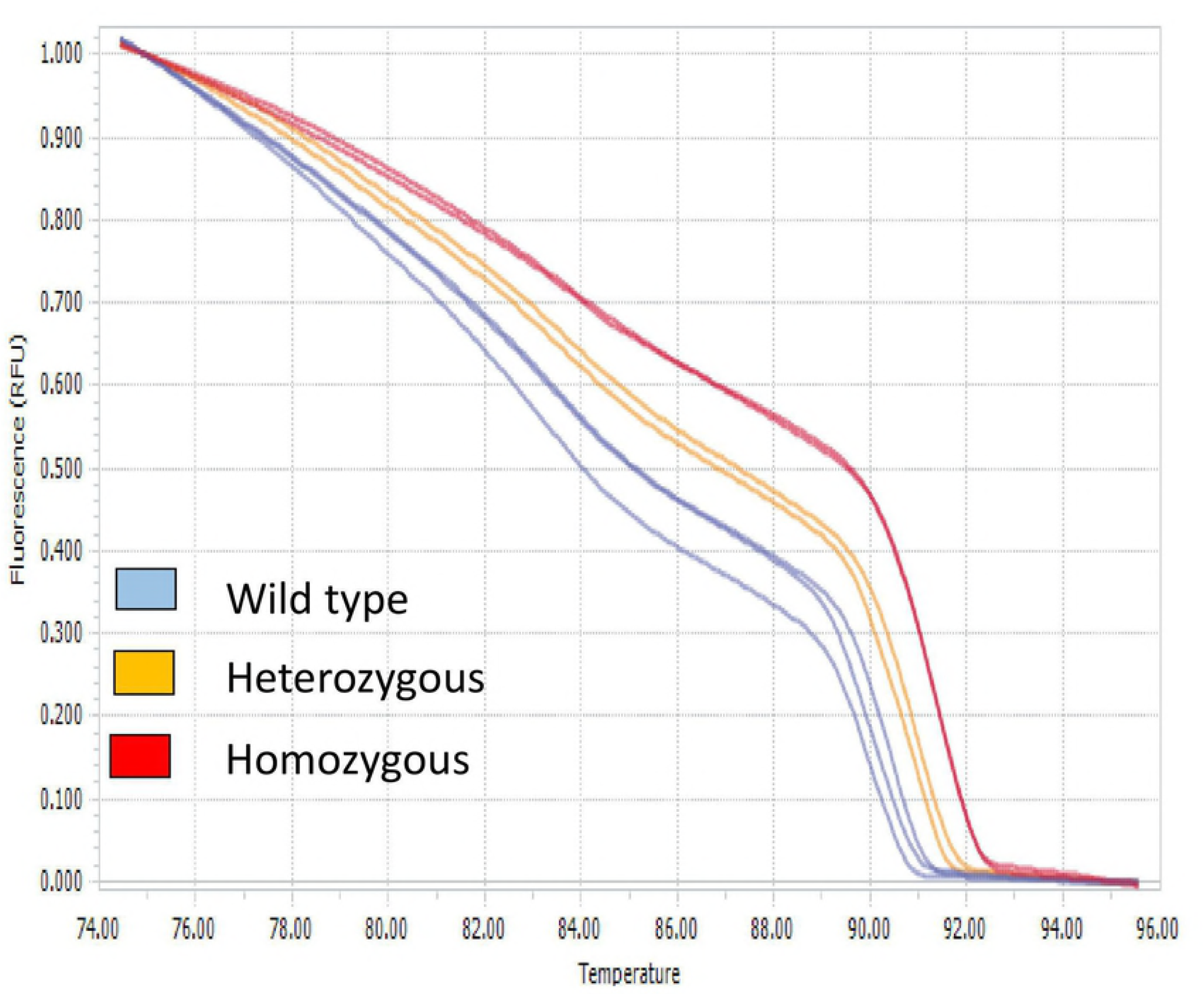
qRT-PCR-HRMA normalized curve for *HSP*-70 Gene

**Fig 2.**
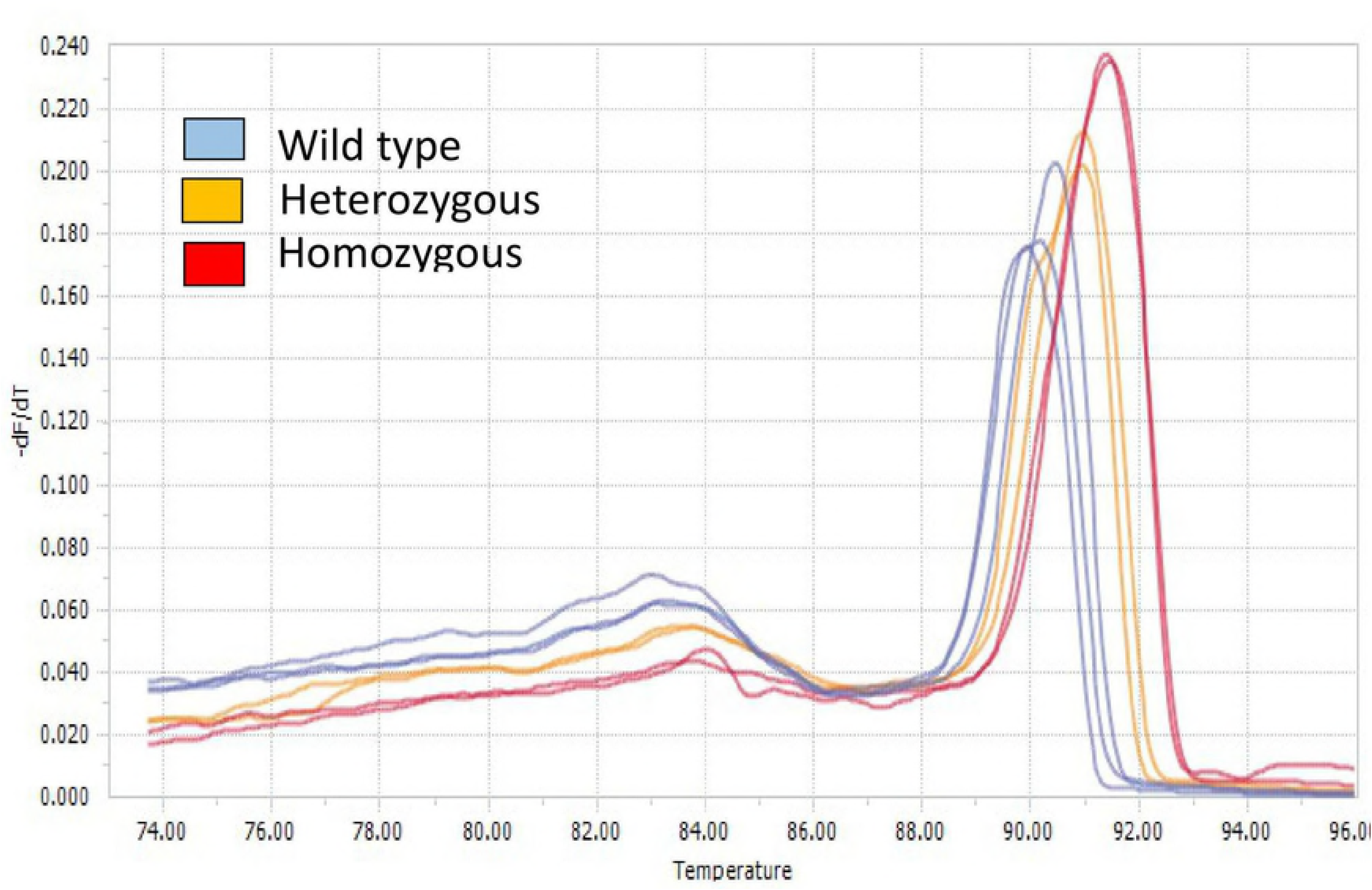
qRT-PCR-HRMA dissociation/derivative curve profile for *HSP*-70 Gene

**Fig 3.**
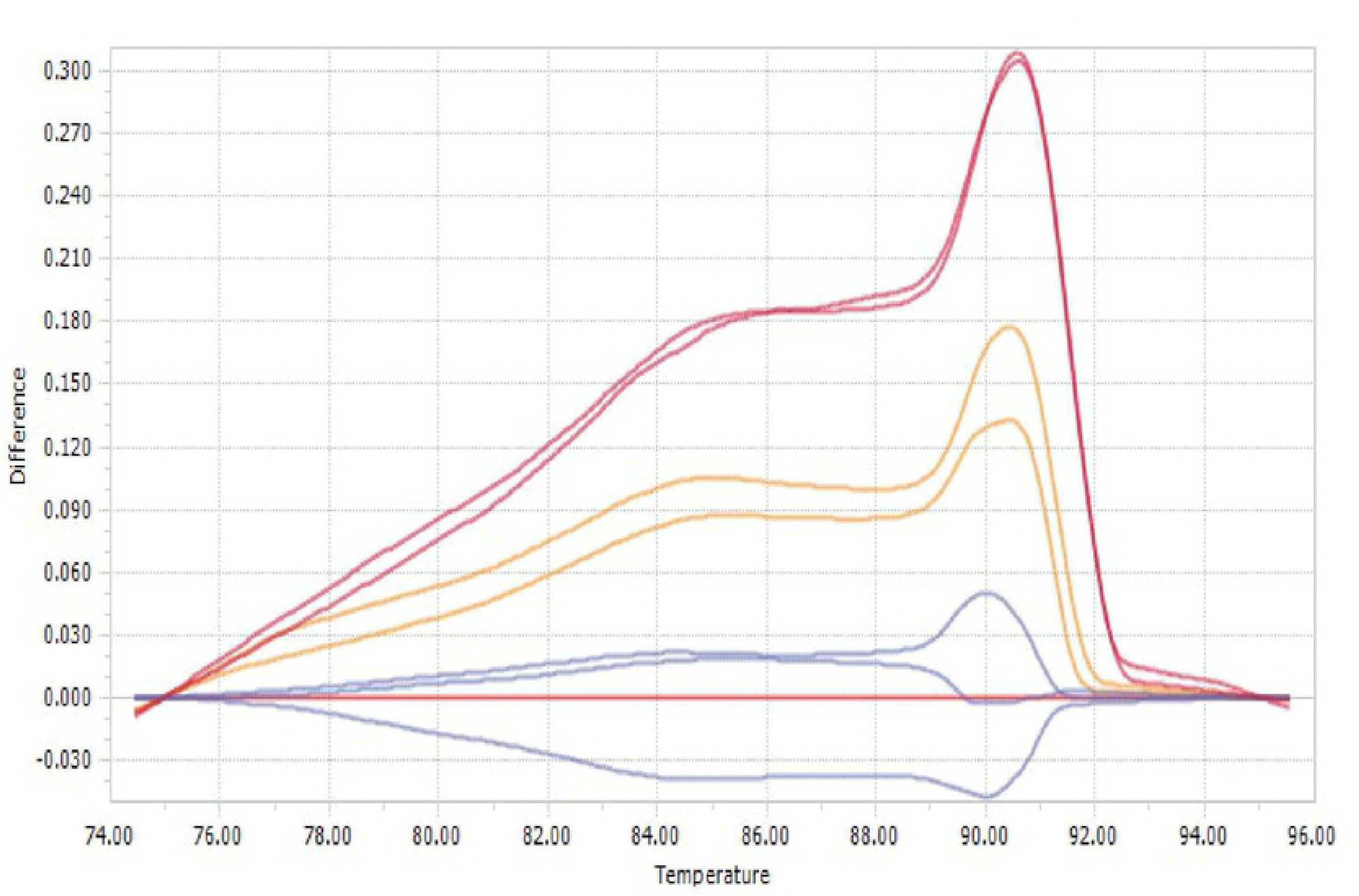
qRT-PCR-HRMA Differential curve for *HSP*-70 Gene

We also detected dissociation of the *HSP*-70 gene sequences into 12 distinct genetic groups at delta ™ discrimination/dissociation (50%) and curve shape discrimination (50%), with fluorescence normalization sensitivity at pre-melting temperature (66.366 °C to 67.366 °C) and post melting temperature range was 94.590 °C to 95.590 °C (Fig. 4 and 5). High Resolution Melting Analysis (HRMA) analysis is a new, post-PCR analysis method used for identifying genetic variation in nucleic acid sequences. It is based on PCR melting (dissociation) curve techniques, enabled by the recent availability of improved double-stranded DNA (dsDNA)–binding dyes along with next-generation real-time PCR instrumentation and analysis software. The method can discriminate DNA sequences based on their composition, length, GC content, or strand complementarity [18]. For the detection of genetic variants, amplicons of 100-300 bp are usually recommended. In this study, variant detection was accomplished with a combination of qRT-PCR and HRMA.

**Fig 4.**
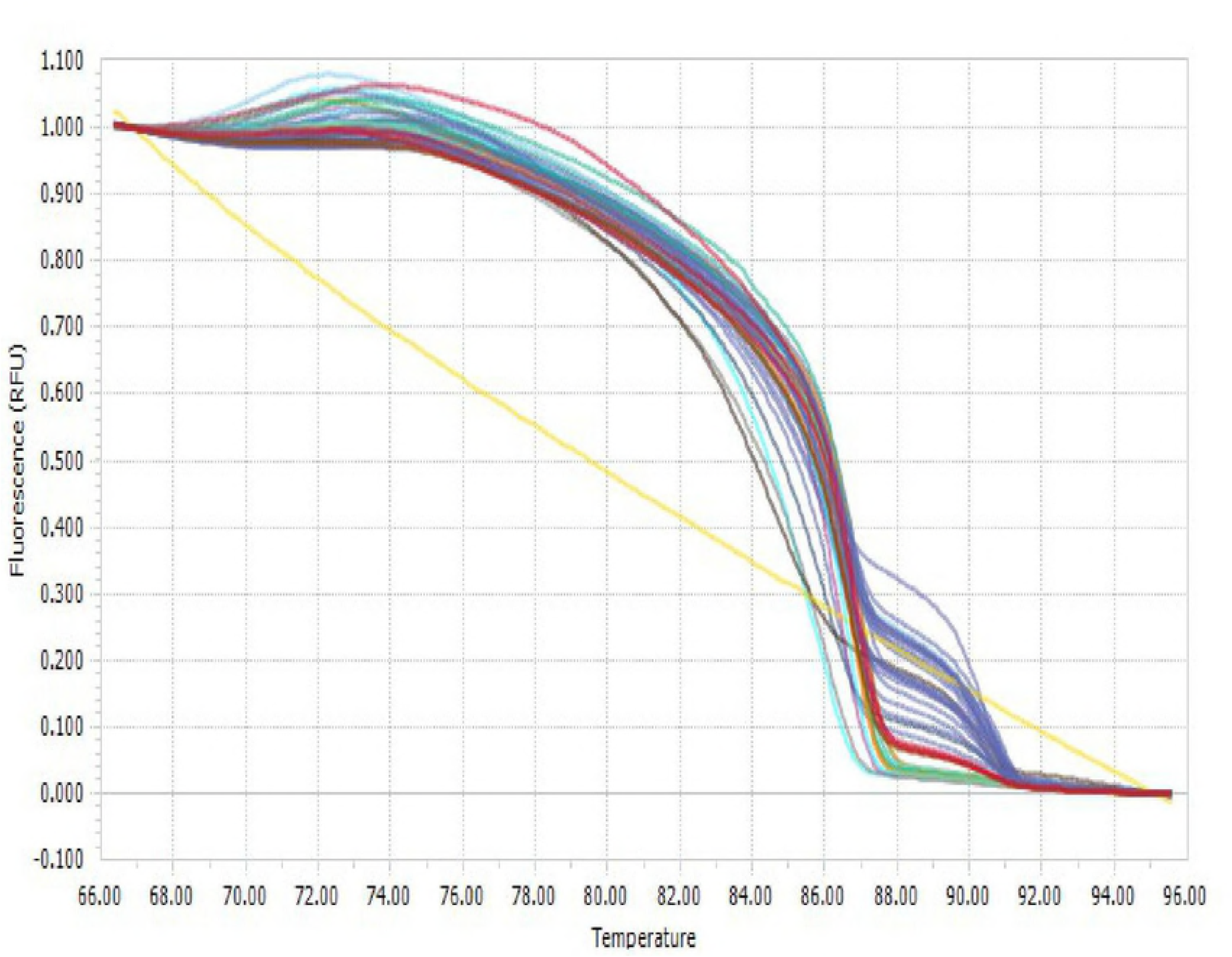
qRT-PCR/HRMA Normalized curve for *HSp-70* gene multiplex of four Nigerian cattle breeds.

**Fig 5.**
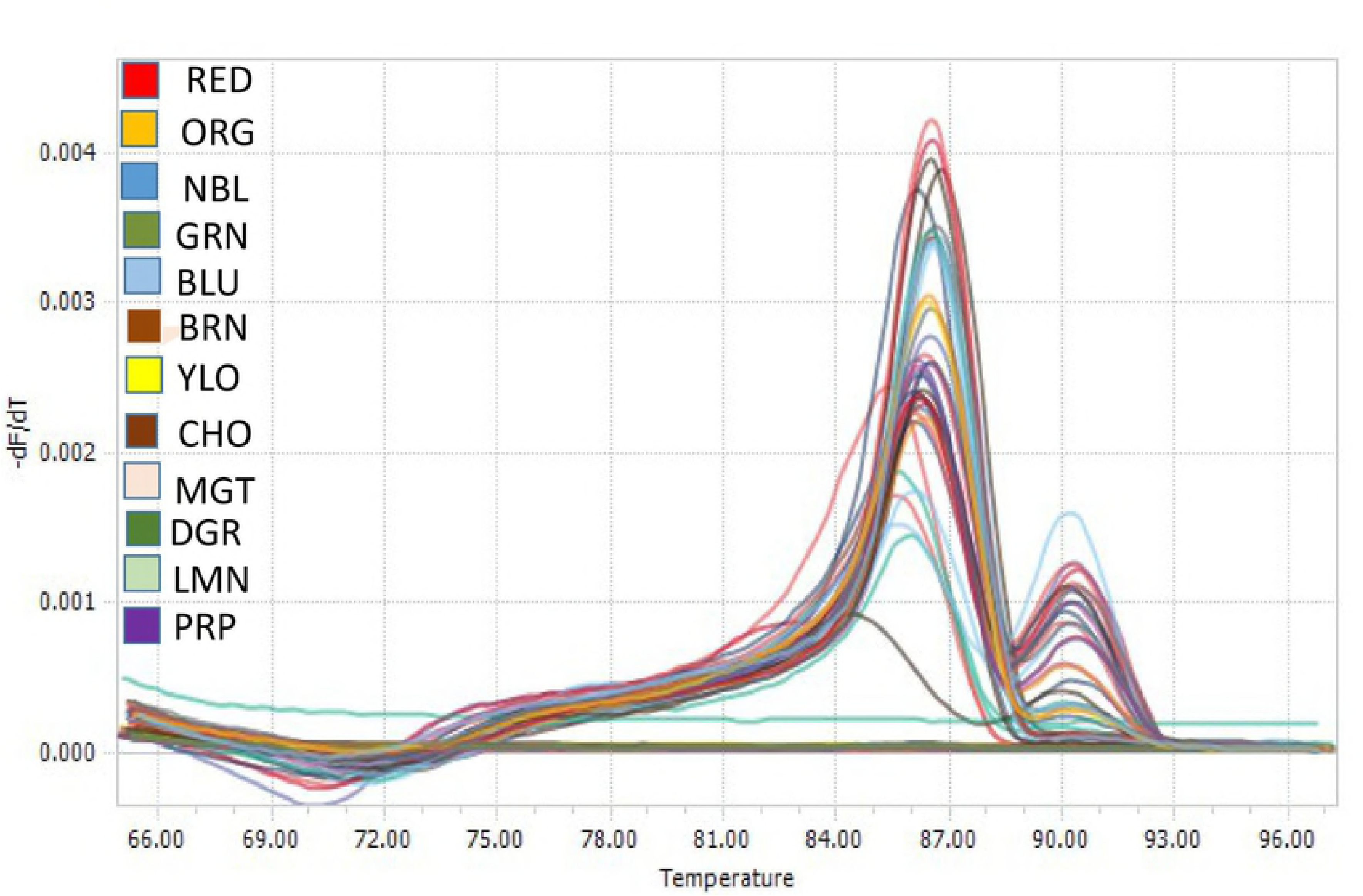
qRT-PCR /HRMA Derivative/ Dissociation curve profile of *HSp-70* gene from four Nigerian cattle breeds.

To detect polymorphism of the gene in the different Zebu cattle breeds under study, we multiplexed and considered normalized qRT-PCR (Figure 4) and normalized HRMA (Fig. 5) separately, then afterwards differential qRT-PCR/HRMA (Fig. 6) together. qRT-PCR/HRMA polymorphisms detection produced 12 distinct genetic variants as depicted with distinct SYBR green fluorescence dye by high resolution melting curve profiles (Table 2). qRT-PCR/HRMA of *HSP-70* gene produced different florescence at a temperature range of 83.54 °C to 86.94 °C (Table 2). Each florescence was associated with gene variant/allele of the gene, which could be the result of single nucleotide difference or nucleotide complexities in the gene sequences from the different cattle breeds. Further analysis by breed showed that gene variants 5 and 8 were common to all four breeds studied, followed by gene variant 7; that was recorded in three breeds. Gene variant 3 was observed in two breeds and all others were observed singly in different breeds. White Fulani had four (4) gene variants. Ambala had seven (7). Genetic diversity detected among the genetic groups (Fig. 4, 5 and 6) suggested that *HSP-70* gene diversity was observed across all the four breeds study. Highest diversity was observed in Ambala cattle population with seven (7) different variants, while the lowest was observed in White Fulani cattle breed with four (4) variants. Sokoto Gudali and Red Bororo had five (5) and six (6) variants respectively. The finding from this study confirms the earlier works of [29] who detected 11 genetic variants in the *HSP-70* gene of Chinese Holstein cattle breed. The result of this study agrees with the reports of [30] who detected polymorphism of *HSP-70* gene in broiler chickens and [31] who reported polymorphism of the *HSP*-*70* gene in humans. This finding is corroborated by the previous works of [32, 33] who reported genetic diversity of *HSP-70* gene in pigs.

**Table 2.**
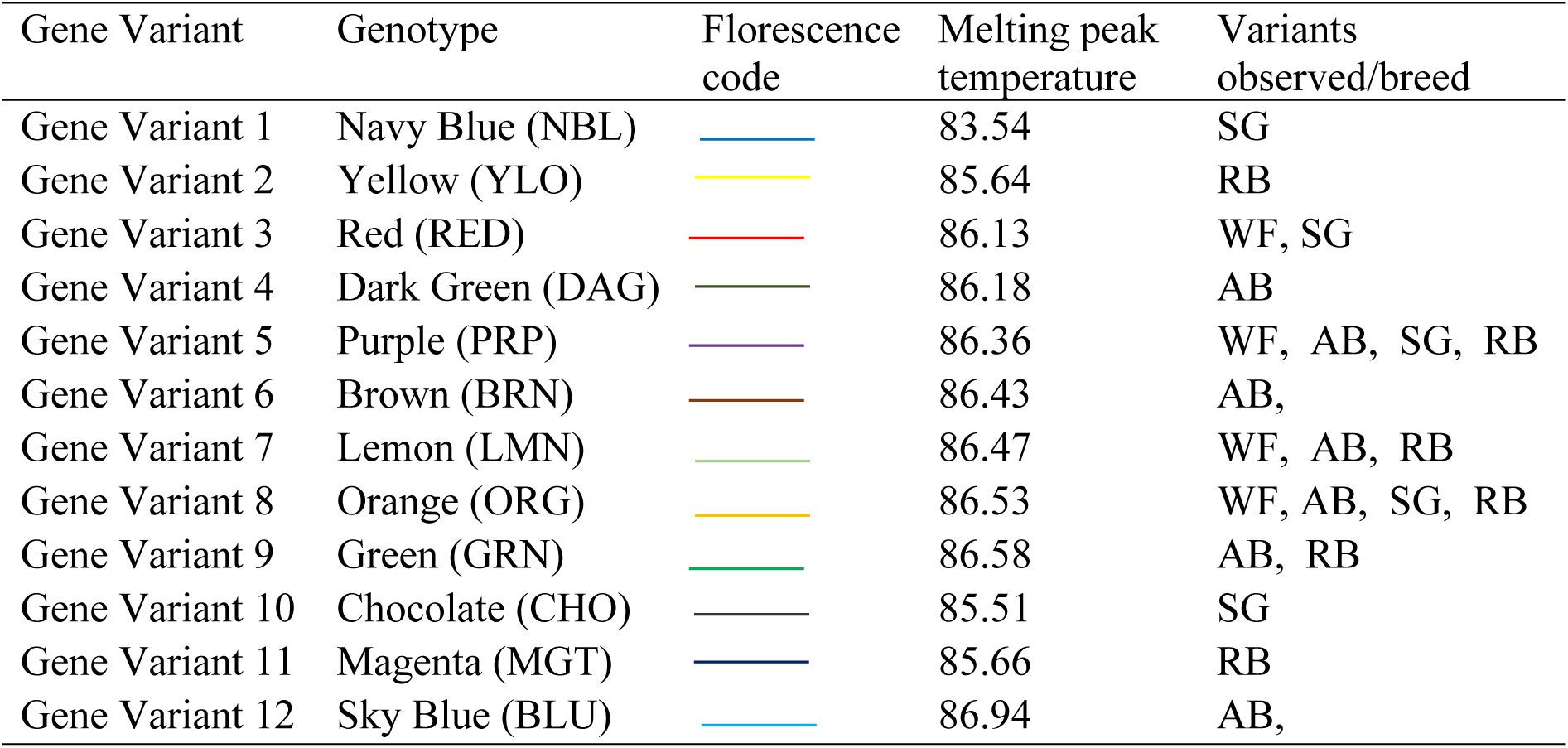
HSP-70 gene variants/alleles distribution in Zebu cattle at different melting peak

| Gene Variant | Genotype | Florescence code | Melting peak temperature | Variants observed/breed |
| --- | --- | --- | --- | --- |
| Gene Variant 1 | Navy Blue (NBL) |  | 83.54 | SG |
| Gene Variant 2 | Yellow (YLO) |  | 85.64 | RB |
| Gene Variant 3 | Red (RED) |  | 86.13 | WF, SG |
| Gene Variant 4 | Dark Green (DAG) |  | 86.18 | AB |
| Gene Variant 5 | Purple (PRP) |  | 86.36 | WF, AB, SG, RB |
| Gene Variant 6 | Brown (BRN) |  | 86.43 | AB, |
| Gene Variant 7 | Lemon (LMN) |  | 86.47 | WF, AB, RB |
| Gene Variant 8 | Orange (ORG) |  | 86.53 | WF, AB, SG, RB |
| Gene Variant 9 | Green (GRN) |  | 86.58 | AB, RB |
| Gene Variant 10 | Chocolate (CHO) |  | 85.51 | SG |
| Gene Variant 11 | Magenta (MGT) |  | 85.66 | RB |
| Gene Variant 12 | Sky Blue (BLU) |  | 86.94 | AB, |

**Fig 6.**
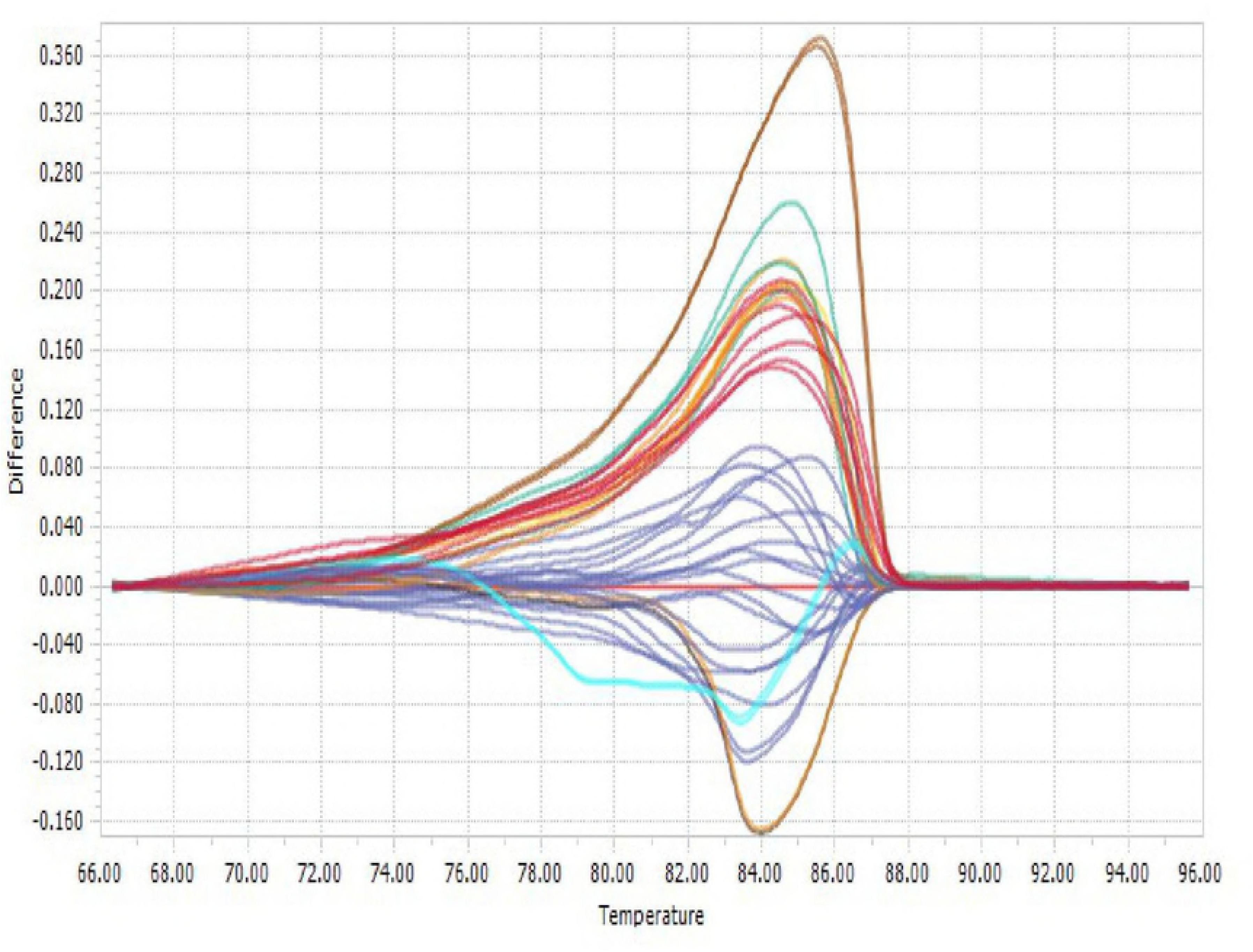
qRT-PCR/HRMA differential plot curve for *HSp-70* gene multiplex in four Nigerian cattle.

qReal-Time PCR/HRMA-based assay, especially in the identification of human and animal pathogens [23-25], genotyping of drug-resistant bacteria isolates [34, 35] and human genetic variants linked to cancer [36, 37] are common. The present study successfully utilized qRT-PCR / HRMA-based assay for polymorphism study of *HSP-70* gene in four Nigerian Zebu breeds of cattle. The findings of this study showed that 12 genetic variants of *HSP-70* gene were detected in four Nigerian zebu cattle studied and agrees with other variant detection studies [20-22].

In this study, we clearly confirmed polymorphism of *HSP-70* gene in Zebu cattle. Polymorphism of the gene in the different breed reflects diversity of the gene in the populations studied. Evolutionary process and diversity of the gene [40,41] is not clear but needs to be investigated. However the present study clearly demonstrated that Nigeria Zebu cattle possess a diversity of *HSP-70*; diversity or heterosis influences evolution and adaptability. Diversity of *HSP-70* gene in Zebu population provides, potentially provides the premise for heat tolerance in the population. Hansen [28] argued that during separate evolution from *Bos taurus*, zebu cattle (*Bos indicus*) had acquired genes that confer thermo-tolerance at the physiological and cellular levels on them. *HSp-70* gene is one specific genes responsible for thermo-tolerance in zebu that have been identified or mapped, and can be put to use in breeding strategies for marker assisted selection for improvement of cattle for heat-tolerance. Adamowicz’s team [38] found a novel single-nucleotide polymorphism (SNP) in the 3’ untranslated region of *HSP-70* gene [29] found five novel mutations in *HSP-70* gene and reported that some of the genotypes confer better thermos-tolerance. Lamb and co-workers [39] also reported eight (8) SNPs in *HSP-70* gene of different cattle breeds and deduced that 5 of them were related to Brahman ancestry.

Past studies by [16, 17, and 29] found variants and mutations in *HSP-70* gene in Holstein cattle that showed differences in thermo-tolerance, but at different loci. These support the findings of the present study that the gene is important for thermo-tolerance in cattle, and further studies to elucidate specific loci associated with thermoregulation and tolerance will be of great use for marker assisted selection and quantitative trait loci (QTL) studies for heat tolerance in cattle. However, the present study is limited in that, the technique used was not designed to pick out distinct variation source and their association with thermo-tolerance traits. Further study that clearly reveals specific single nucleotide polymorphisms (SNPs) that can be explored for association with heat tolerance trait is quite important.

## Conclusion

Heat stress is one of the critical factors among various environmental stressors that impede profitable livestock rearing. Our results elucidated 12 genetic variants of *HSP-70* gene in four Nigerian zebu cattle detected through multiplex qRT-PCR-HRMA based assay. We therefore hypothesise that the detected novel genetic variants of *HSP-70* gene of Nigerian zebu cattle could be interrogated as veritable genetic resource for improvement programme for selection of thermo-tolerance ability, adaptability and survivability advantage to cope with wide range of thermal stress and environmental variations especially in the hot humid tropics. The study supports qRT-PCR-HRMA genotyping for gene characterization. However, the limitation of the technique is found in its inability to generate specific SNPs and the gene loci they reside. Further studies to elucidate specific loci associated with thermoregulation and tolerance will be of great use for marker assisted selection and quantitative trait loci (QTL) studies for heat tolerance in cattle.

## Acknowledgements

The Government of India funded the main analyses of this study in a form of visiting scholarships through the Research Fellowship for Developing Countries Scientists (RFD-CS). This fellowship is administered by the Department of Science and Technology (DST) and Ministry of External Affairs (MEA) of the Government of India, New Delhi. We thank the management of slaughter houses in Nigeria for giving us permission to sample from the animals in their custody.

